# Time efficient preparation methods for MRI brain scanning in awake young children and factors associated with success

**DOI:** 10.1101/259358

**Authors:** Camilia Thieba, Ashleigh Frayne, Matthew Walton, Alyssa Mah, Alina Benischek, Deborah Dewey, Catherine Lebel

## Abstract

**Objective:** Young children are often unable to remain still for magnetic resonance imaging (MRI). Various preparation methods have been reported to avoid sedation or anesthesia, with mixed success rates and feasibility. Here we describe a time-efficient preparation method and factors associated with successful scanning in young chdilren.

We recruited 134 children aged 2.0–5.0 years for an MRI study. Some children completed a training session on a mock scanner, and all children received a 15–20 minute introduction to scanning procedures immediately before their scan. We compared success between children receiving mock scanner training or not, and evaluated demographic or cognitive factors that predicted success.

**Results:** 97 children (72%) completed at least one sequence successfully; 64 children provided high-quality data for all 3 sequences. Cognitive scores were higher in successful children, but children who received mock scanner training were less likely to be successful. A case-controlled comparison of children matched on age, gender, and cognitive scores found no differences between children receiving training or not.

We present a quick method for preparing young children for awake MRI scans. Our data suggests limited advantages of mock scanner preparation for healthy young children, and that cognitive abilities may help predict success.

## Introduction

Magnetic resonance imaging (MRI) is a non-invasive technique useful for numerous research and clinical applications, though it is very sensitive to motion. Collecting high quality data sets is particularly difficult during the preschool years (~2–4 years), as children do not readily fall asleep or follow instructions to stay still. Sedation or general anaesthesia are often used in clinical settings [1, 2], but are not appropriate for research [1, 3–5]. Scanning children during natural sleep can be quite successful [4, 6–8], but requires flexible scanning schedules (i.e., evening/night time), and may involve long time periods waiting for children to fall asleep. Preparation techniques for scanning awake children include tours of the MRI facility, training in a mock scanner, and play therapy [6, 9–14]; an audio/visual system can also increase compliance [3, 13, 15, 16]. A mock scanner training protocol resulted in 72% and 54% success for children 3.7–4.9 years for structural and functional scanning, respectively [10]; other studies reported 66% success in children 0–4.5 years [6] and 97% success for 45 children aged 4–6 years [14]. In a clinical MRI study, sedation rates were reduced from 89% to 67% in children 2–4 years, equivalent to 33% success [15].

In our ongoing study of brain development in young children, we use a rocketship themed training protocol, with optional mock scanner training. Here, we describe the protocol and evaluate factors associated with children’s success. This protocol requires little preparation time and can be implemented in centres lacking flexible scanning hours.

## Main Text

### Materials and Methods

#### Participants

This was a retrospective analysis of data collected for a different study examining brain development in early childhood [17]. 134 children aged 1.96 – 4.95 years (3.4 +/− 0.6 years) were recruited to that study. All were English speakers born full term, and free from genetic disorders associated with significant intellectual or motor impairments, neurological or neurodevelopmental disorders, and contraindications to MRI. Written informed consent was obtained from a parent/guardian. The Conjoint Health Research Ethics Board at the University of Calgary approved this study, REB13-0020.

At the time of the MRI scan, years of maternal post-secondary education was collected, as well as two language measures on all children aged 3 years or older: the Phonological Processing and Speeded Naming from the NEPSY-II [18]. Some children were part of another (non-imaging) study [19] and were assessed approximately 1 year prior to scanning on the Cognitive, Language, and Motor Composite scales of the Bayley Scales of Infant and Toddler Development-III (Bayley-III) [20] (n=104) and the Attention, Internalizing, and Externalizing Behavior Problems scales of the Achenbach System of Empirically Based Assessment [21] (ASEBA; n=96). See Table 1.

**Table 1:**
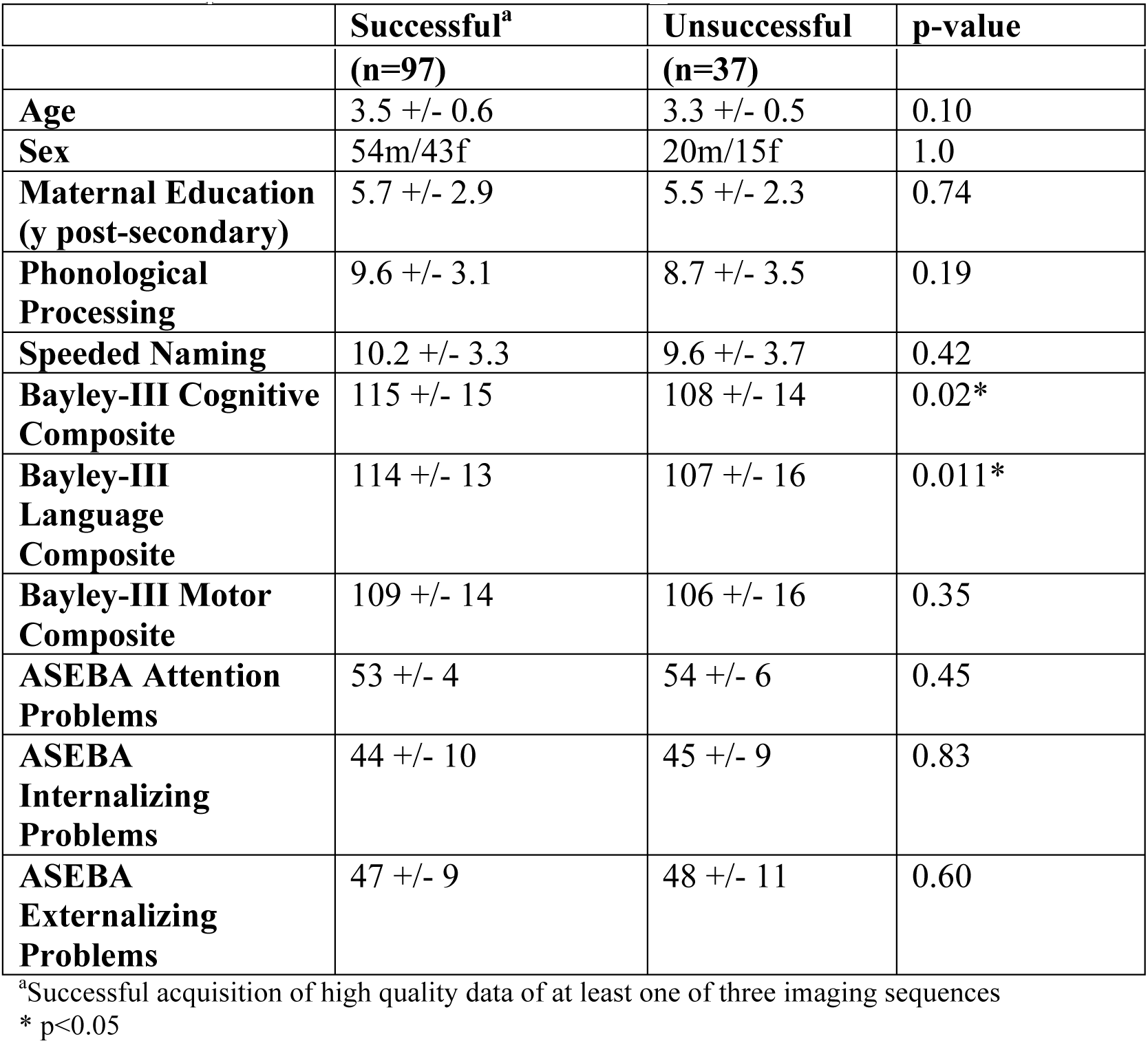
Demographic characteristics of children participating in the study. Groups are separated based on scan success, with success defined as at least one sequence with high-quality data. Two-sample t-tests were used to test for group differences.

#### Pre-Scanning Training Sessions

At the time of enrollement, parents were given audio, video, and website resources about MRI scanning, and encouraged to discuss the procedures with their child. These resources included a link to download our e-book, *Pluto and the MRI Rocket Ship Adventure* (Figure 1; available for free download: https://www.lulu.com/shop/search.ep?contributorId=1347527).

**Figure 1.**
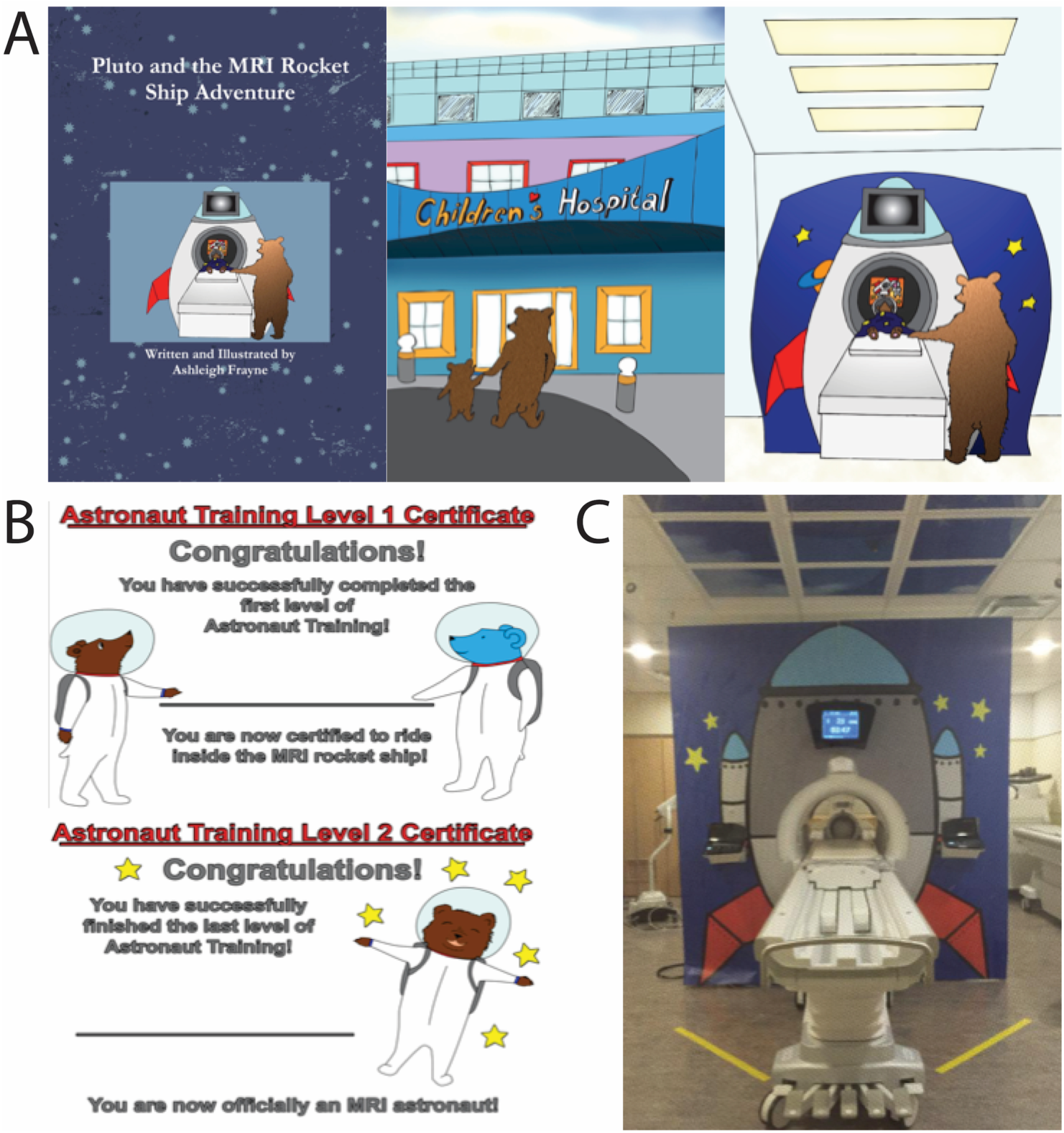
Space adventure-themed training materials. Space adventure-themed training materials include a children’s book about an MRI Rocket ship adventure (A), astronaut training certificates (B), and a rocket ship façade for our MRI scanner (C).

Our facility has a mock scanner equipped with a rocket ship façade identical to that on the real MRI scanner and the MRI rocketship in our book (Fig. 1). Families were given the choice to participate in mock MRI scanner training or not; 15% (n=20) chose to complete a training session. Common reasons for opting out were time constraints or because parents thought their child “did not need it”. Mock scanner training occured within 7 days of the MRI scan, and consisted of briefly describing the MRI procedure, introducing the child to the scanner, putting the parent or a stuffed animal into the bore, and having the child practice lying still within the bore while watching a movie and listening to typical MRI scan sounds. At the end of the training, children were given an astronaut training certificate. Training sessions lasted approximately 30 minutes.

All families were contacted again 1–2 days prior to the real MRI scan and reminded to review the preparation materials, and bring a favorite movie and/or comfort item to the scan. On the day of the scan, children were met by a research assistant. The research assistant and the child then played short games to get comfortable, completed a 10-minute language assessment, read the rocketship story, and went over MRI scan steps and expectations. The child was told about the “big camera” that makes loud noises, instructed to lie still while “pictures” were being taken, given stickers and promised a gift at the end of the procedure. A teddy bear was given to each child to take with them inside the scanner, as described in our book. The family was taken to the scanner and given the opportunity to quickly familiarize themselves with the environment (~5 minutes). The child was shown the equipment and asked to lie down on the scanner bed, where he/she was fitted with headphones and positioned inside the head coil, then inside the bore. During scanning, a parent remained beside the child, and children watched movies on the projector screen. Children were reminded to “hold still like a statue”. Total time spent prior to imaging, including the 10-minute language assessment, was approximately 30 minutes; only 5 minutes of this preparation time was spent in the MRI scanning environment. At the completion of the scan, children were given an astronaut certificate for their “space flight”, and a toy.

#### MR Imaging

Imaging was performed during daytime or early evening hours (<7 pm) at the Alberta Children’s Hospital on the research-dedicated 3T GE MR750w MRI scanner (General Electric, Waukesha, WI) with a 32-channel head coil. The protocol consisted of (in order): diffusion tensor imaging (DTI; 4:03 min:s), anatomical T1-weighted imaging (4:12), and T2*-weighted imaging (4:12) (Table S1). Additional sequences, including arterial spin labeling, single voxel spectroscopy, and resting state functional MRI were acquired on participants who were comfortable when time permitted; these are not discussed here. Foam padding was used to minimize head motion.

#### Scan Quality Assessment

In-house Matlab software was used to detect DTI volumes with excessive motion, which were removed prior to data analysis. Scans with > 22 volumes (66%) retained were suitable for tractography and voxel-based analysis, and considered successful. T1-weighted image quality was rated on a five-point scale, where 1 represented unusable data and 5 represented no motion artefacts (Figure S1). Scans with scores of 3–5 were considered successful and of sufficiently high quality for processing through FreeSurfer [22]. T2*-weighted images were assessed using a similar rating scale, where scores of 3–5 were considered successful and were used in further analysis (Figure S1).

#### Statistical Methods

Statistical analysis was performed in SPSS version 24.0. Two-sample t-tests were used to investigate demographic, cognitive, and behavioral differences between children who provided good quality data and those who did not. Demographics and quality measures (DTI volumes retained, T1 score, T2* score) were also compared between children who received mock scanner training and those who did not, using non-parametric tests for two independent samples. The association between mock scanner training and scan success was tested using a chi-squared test.

## Results

### Image Quality

We report results here for the first attempt at scanning (i.e., no one returned for a second visit). Median ratings for T1-weighted and T2*-weighted images were 3; the mean number of usable DTI volumes was 31 (of 35) (Figure S2). Sixty children (48%) provided high quality scans for all sequences (T1, T2*, DTI). An additional 10% provided high quality data for 2 sequences and 14% provided high quality data for only 1 sequence. The 96 children (72%) with at least one high quality dataset were termed the “successful” group, and the other 37 (28%) termed the “unsuccessful” group.

### Scanning Success

The successful group had significantly higher Bayley-III Cognitive and Language Composite scores than the unsuccessful group; no significant differences were found for maternal education, child’s age, sex, or behavior (Table 1).

### Training differences

Mock scanner training was not associated with scan success, but there were trends where children receiving training were less likely to be successful overall (p=0.1), and less likely to be successful on T1 imaging (p=0.059) than children with no mock scanner training (Table 2). Children who completed training also had significantly lower Bayley-III Cognitive and Language and NEPSY-II Phonological Processing scores, and higher quality ratings on T1-weighted images (Table 2).

**Table 2:**
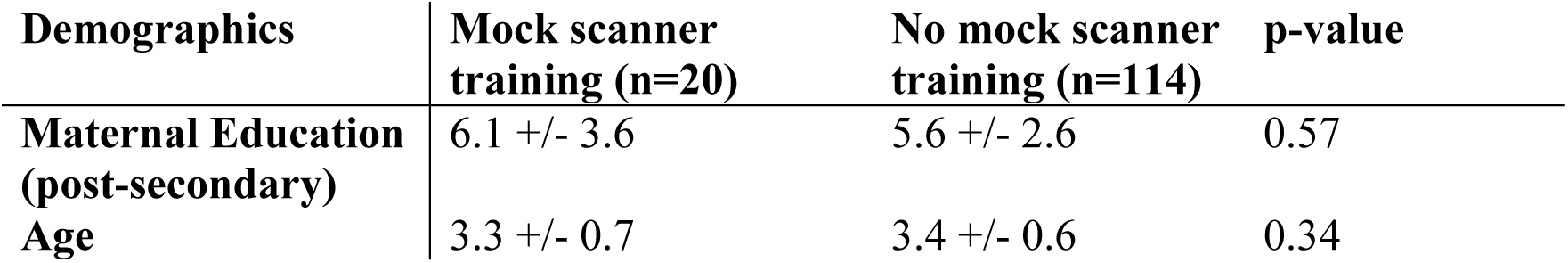
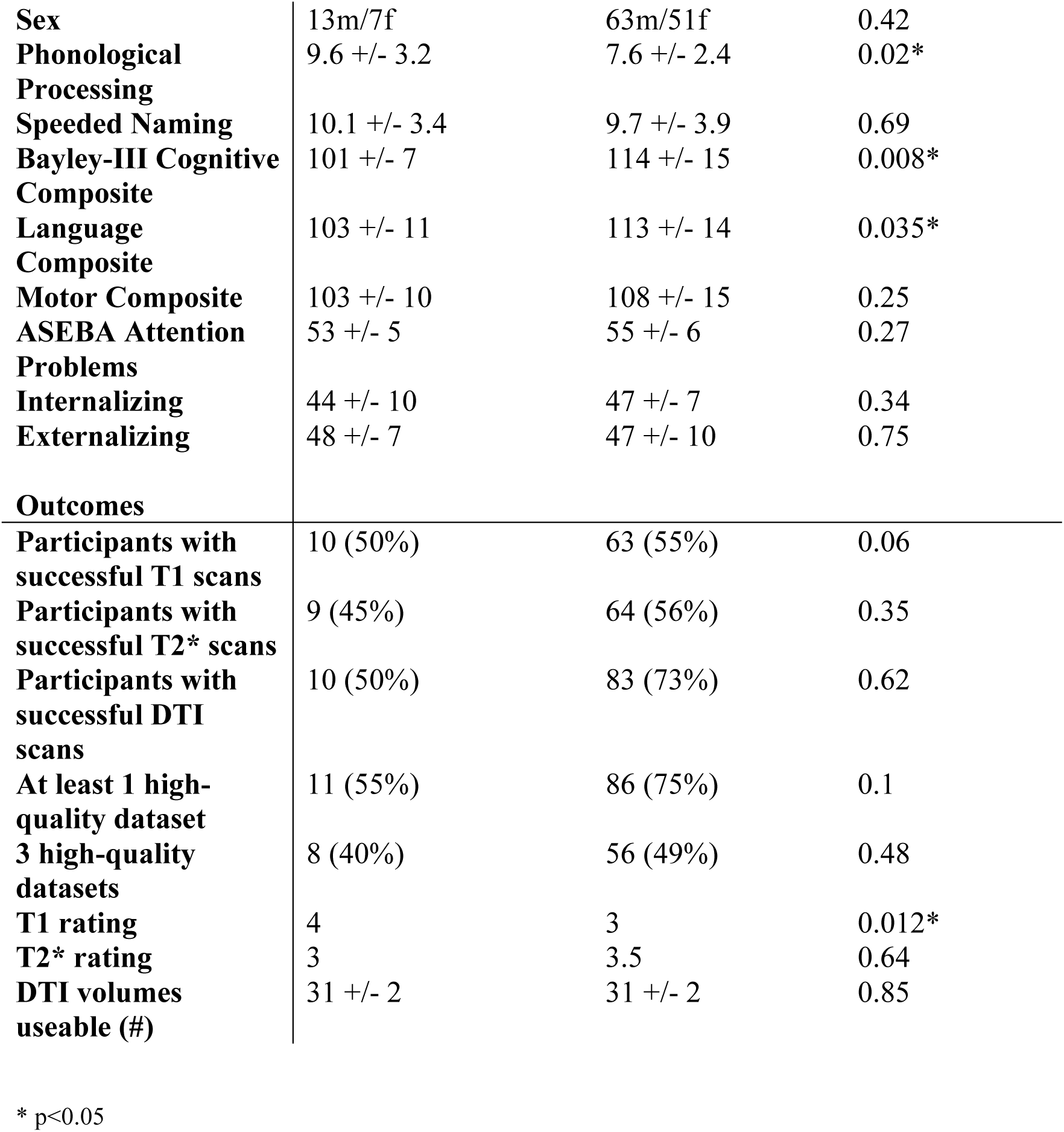
Differences between groups receiving mock scanner training or not. Two-sample t-tests were used to test for group differences; non-parametric tests were used for image quality ratings.

### Case-control comparison analysis

Because of the apparent sample bias in the mock scanner training group (lower cognitive and language scores), a case-control analysis was conducted that matched children in the two groups on Phonological Processing scores, age and sex. Only 17 mock scanner participants had Phonological Processing scores, so each matched group contained 17 participants. Results revealed no significant group differences on cognition, behaviour, or scan outcomes (Table S2).

## Discussion

Successful pediatric neuroimaging sessions are important for studying early childhood brain development, and for clinical assessments. Here, we describe a time-efficient method for preparing young children for MRI scans that resulted in high success rates (72%) on the first scan attempt. Our preparation method took approximately 15–20 minutes immediately prior to the MRI scan (i.e., no additional visit), with most preparation occurring outside the scanning environment. This methodology could be easily applied in an environment where scanning children during natural sleep is difficult due to time or scheduling constraints.

While the idea of using storytelling and imagination to engage children in scanning sessions is not unique [11], customizing training tools to engage children with a specific location and research group is a fairly new concept. Our training tools featured key elements of our scanning site and introduced children to the research staff involved in the training sessions. This technique helped to establish rapport with children and families prior to the onsite visit, possibly shortening the time required to build children’s trust.

Few studies report success rates specifically in preschool-aged children. One study had a 54–71% success rate in 4 year-olds [10], and another reported 33% success for clinical scans in awake 2–4 year olds [15], comparable to or lower than our success rate of 72%. Success rates across wider age ranges are 66% [6] to 97% [7, 14], and tend to be higher in studies that require advance training visits, multiple attempts at scanning, and/or preparation times over 1 hour [6–8, 14]. Therefore, while more preparation may be beneficial, our results indicate that scanning of young children is also possible with short preparation times.

Children in the successful group had higher cognitive scores than children in the unsuccessful group, suggesting that cognition may help predict success during scanning. Furthermore, parents may be good predictors of their children’s success, since the self-selected group with no mock scanner training was more successful overall. A case-control analysis in 34 children matched for age, gender, and Phonological Processing scores revealed no significant differences between training groups on any variables. This contrasts with previous findings in adults that showed mock scanner training to be effective in reducing subjective distress [23], and suggests that young children may not benefit from training as much as older children or adults.

In conclusion, our results suggest that cognitive scores or parents may help predict young children’s success in MRI scanning, but our limited data does not show a substantial advantage of mock scanner training. Ultimately, this study shows the feasibility of conducting MRI exams in awake preschool-aged children, and demonstrates a need for further research regarding training protocols and variables that predict success.

## Limitations

The main limitation of this study is that it was not a randomized trial of training protocols, and there was a sample bias among children who had mock scanner training. When a case-controlled analysis was performed, only 17 children could be matched on cognitive/language scores, resulting in a small sample size. Future research using randomized designs and larger samples is necessary to ascertain the true effects of training protocols and other factors on scan success in children. Another limitation is that we did not measure to what extent families used the preparation materials, and this likely varied considerably. Future studies collecting this information would be valuable.

### List of abbreviations

MRI: magnetic resonance imaging. ASEBA: Achenbach System of Empirically Based Assessment. DTI: diffusion tensor imaging.

## Declarations

### Ethics approval and consent to participate

This study was approved by the University of Calgary conjoint health research ethics board; approval number REB13-0020. All participants’ parent or guardian provided written informed consent. Children provided verbal assent to participate.

### Consent for publication

Not applicable.

### Availability of data and material

The datasets generated and/or analyzed during the current study are available from the corresponding author on reasonable request.

### Funding

This work was supported by the Canadian Institutes of Health Research (CIHR) (funding reference numbers IHD-134090, MOP-136797, and a New Investigator award to CL), and a grant from the Alberta Children’s Hospital Foundation.

Authors’ contributions. CT, AF, and CL performed data analysis and interpretation. CT, AM, MW, and AB performed image analysis and quality assessment. AF, DD, and CL wrote the manuscript. All authors read and approved the final manuscript.

## Acknowledgments

The authors thank the members of the APrON study team for recruitment assistance.

